# Using Conditional Generative Adversarial Networks to Boost the Performance of Machine Learning in Microbiome Datasets

**DOI:** 10.1101/2020.05.18.102814

**Authors:** Derek Reiman, Yang Dai

## Abstract

The microbiome of the human body has been shown to have profound effects on physiological regulation and disease pathogenesis. However, association analysis based on statistical modeling of microbiome data has continued to be a challenge due to inherent noise, complexity of the data, and high cost of collecting large number of samples. To address this challenge, we employed a deep learning framework to construct a data-driven simulation of microbiome data using a conditional generative adversarial network. Conditional generative adversarial networks train two models against each other while leveraging side information learn from a given dataset to compute larger simulated datasets that are representative of the original dataset. In our study, we used a cohorts of patients with inflammatory bowel disease to show that not only can the generative adversarial network generate samples representative of the original data based on multiple diversity metrics, but also that training machine learning models on the synthetic samples can improve disease prediction through data augmentation. In addition, we also show that the synthetic samples generated by this cohort can boost disease prediction of a different external cohort.

## 1 INTRODUCTION

The microbiome is a collection of microscopic organisms cohabitating in a single environment. These organisms have been shown to have a profound impact on its environment. Of particular interest is the human microbiome and how its composition can affect the health and development of the host. In particular, the microbiome of the human gut has been linked to the pathogenesis of metabolic diseases such as obesity, diabetes mellitus, and inflammatory bowel disease (Barlow, Yu, & Mathur, 2015; Franzosa et al., 2019; Tilg & Kaser, 2011). Additionally, the gut microbiome has been shown to have an effect on the development and modulation of the central nervous system (Carabotti, Scirocco, Maselli, & Severi, 2015), stimulation of the immune system (Fung, Olson, & Hsiao, 2017), and even impact the response to cancer immunotherapy treatment (Gopalakrishnan, Helmink, Spencer, Reuben, & Wargo, 2018). Because of the profound effect that the microbiome has on the human host, it is of increasing importance to understand how the changes in its composition lead to physiological changes in the host.

An important analysis in microbiome studies involves uncovering underlying association between microbes and the host’s health status. However, modelling the underlying distribution of microbiome data has been a long-standing challenge due to the sparsity and over-dispersion found in microbiome data. There have been many statistical approaches proposed for modelling microbiome data over the past decade, however there is still no consensus as to which models and underlying assumptions are best suited for handling the complexity of the data. (Kurilshikov, Wijmenga, Fu, & Zhernakova, 2017; Xu, Paterson, Turpin, & Xu, 2015).

Recently, machine learning (ML) models have been advocated for a data-driven approach for the prediction of the host phenotype (Knights, Parfrey, Zaneveld, Lozupone, & Knight, 2011; LaPierre, Ju, Zhou, & Wang, 2019; Pasolli, Truong, Malik, Waldron, & Segata, 2016). However, one persistent challenge is the relatively small sample size in microbiome datasets. It is often the case that microbiome datasets have a far greater number of features than the number of samples, which can quickly lead to the overfitting of models.

To address these challenges and limitations, we construct a novel method for generating microbiome data using a conditional generative adversarial network (CGAN). We then construct synthetic samples using the generative model in order to augment the original training set. Data augmentation is a technique often used in ML to improve task performance and improve generalization (Bowles et al., 2018; Mikołajczyk & Grochowski, 2018). By generating a large number of synthetic microbiome samples that resemble the original data, we show that it is possible to improve the performance of ML models trained on the generated synthetic samples.

Generative adversarial networks (GANs) involve two neural networks competing against each other in an adversarial fashion in order to learn a generative model in a non-parametric data-driven approach (Goodfellow et al., 2014). GAN models have shown success in multiple domains including the generation of medical images (Frid-Adar et al., 2018) and single cell RNA-Seq gene expression profiles (Ghahramani, Watt, & Luscombe, 2018). Additionally, synthetic datasets generated using GAN models have shown to be able to boost performance of prediction based tasks through data augmentation (Che, Cheng, Zhai, Sun, & Liu, 2017). A recent study has also explored the behaviour of Wasserstein GAN models with gradient penalty in microbiome data, showing success in generating realistic data compared to other simulation techniques (Rong et al., 2019). However, the authors do not fully explore improvements in analyses using the generated data.

We use a variation of standard GAN models called CGAN. CGANs incorporate side information into the model to allow the generation of samples from different distributions given certain underlying conditions, such as disease status, which has shown improvement from standard GAN models (Mirza & Osindero, 2014).

The main contribution of this manuscript is the utilization of the CGAN model in order to construct a generator that can sample from different conditions to provide synthetic data representative of the true data. Additionally, we use the generator to synthesize samples for data augmentation. We show that the generated data not only are similar to the original data with respect to diversity metrics, but also that the data augmentation can lead to statistically significant improvement in performance of ML models.

## 2 MATERIALS AND METHODS

### 2.1 Datasets used in study

For our study, we use the data reported from two different cohorts of patients with inflammatory bowel disease (IBD). The Prospective Registry in IBD Study at MGH (PRISM) enrolled patients with a diagnosis of IBD based on endoscopic, radiographic, and histological evidence of either Crohn’s Disease or Ulcerative Colitis. The second dataset is used specifically for external validation and consists of two independent cohorts from the Netherlands (Tigchelaar et al., 2015). The first consists of 22 healthy subjects who participated in the general population study LifeLines-DEEP in the northern Netherlands. The second cohort consists of subjects with with IBD from the Department of Gastroenterology and Hepatology, University Medical Center Groningen, Netherlands. This will be used as the validation dataset.

Processing of the stool samples collected for both datasets is described in the original study (Franzosa et al., 2019). Briefly, metagenomic data generation and processing were performed at the Broad Institute in Cambridge, MA. Quality control for raw sequence reads was performed using KneadData v0.5.1. Reads were then taxonomically profiled using MetaPhlAn2 (Segata et al., 2012) and relative abundance was collected at the species level. The relative abundance values are publicly available and were obtained from the original study (Franzosa et al., 2019). A summary showing the number of IBD patients, healthy subjects, and species level microbes for each dataset is shown in Table 1.

**Table 1:** Datasets used in study.

|  | # IBD | # Healthy | # Microbes |
| --- | --- | --- | --- |
| PRISM | 121 | 34 | 195 |
| Validation | 43 | 33 | 115 |

### 2.2 CGAN architecture

In order to generate synthetic microbial community structures, we utilize a CGAN architecture. A standard GAN is composed of two competing networks: a generator and a discriminator. The task of the generator is to learn to generate synthetic data representative of real data while the discriminator tries to predict if a given sample is synthetic or real. The generator is trained to maximize the probability of the discriminator in misclassifying samples. At the same time, the discriminator is trained to minimize this probability. A CGAN expands on standard GAN models by feeding side information, i.e., the disease status, to both the generator and discriminator. This allows the generator to generate synthetic samples conditioned on the provided side information.

The generator, *G*, of the CGAN model requires two sets of inputs: a set of priors and the conditional side information. In our study, we sample our priors from the uniform distribution ∼U(−1,1). Both inputs are fed through multiple fully connected hidden layers of perceptrons and finally to an output layer. The output of the generator represents a vector of microbial abundance features.

The discriminator, *D*, takes a sample of microbial abundance features as an input in addition to the side information. The inputs are passed through multiple fully connected layers and then to an output of a single node using the sigmoid activation function. The sigmoid function is used in order to squish values to be between 0 and 1. The output of the discriminator represents the prediction of the probability that the given sample of data is real.

Both generator and discriminator networks are trained in an iterative fashion such that in each epoch, the discriminator is first trained on the generated and real samples and the weights are updated. After the discriminator has been updated, the generator is updated. The loss functions for the discriminator and generator are shown below.

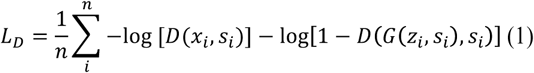

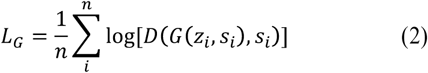

Here *n* represents the number of real samples, *z*_*i*_ represents a vector of priors for the generator, *x*_*i*_ is the relative abundance vector of a real microbial community sample, and *S*_*i*_ is the side information that the networks are conditioned on. *D*(*x*_*i*_, *S*_*i*_) is the discriminator’s prediction if *x*_*i*_ is real given the side information *S*_*i*_. *G*(*z*_*i*_, *S*_*i*_) is the generator’s prediction of a synthetic sample given the prior noise *z*_*i*_ and side information *S*_*i*_ A figure showing the architecture of our CGAN is shown in Fig.1.

**Figure 1:**
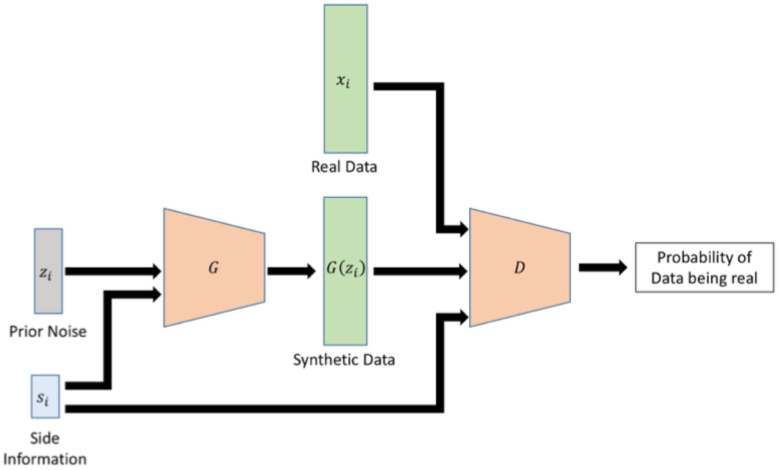
Visualization of the CGAN architecture. A set of prior noise *z*_*i*_ and side information *S*_*i*_ corresponding to sample *x*_*i*_ are used to generate a synthetic sample. The discriminator then uses the side information to predict if a given sample is real or synthetic.

## 3 RESULTS

### 3.1 CGAN training

In our analysis, we sample a vector of size 8 for *z*_*i*_ in the generator model. We add a one-hot encoded value of the disease state (IBD or healthy) as *S*_*i*_ and concatenate the two inputs together. The concatenated input is then passed through two fully connected layers of size 128. Batch normalization is performed at each layer. The leaky ReLU activation function with an alpha value of 0.1 is performed after each batch normalization. Unlike the standard ReLU activation function, leaky ReLU still allows a small positive gradient when given negative values. The output layer of the generator uses a softmax activation function in order to reconstruct the relative abundance of the microbial community.

The discriminator network takes a vector or microbial relative abundance features as well as the one-hot encoded disease state for that sample. The two inputs are concatenated and fed through two fully connected layers of size 128. The leaky ReLU activation is again used for each fully connected layer. The output of the discriminator was a single node with a sigmoid activation to shrink the prediction value to be between 0 and 1.

CGAN models were trained only using the PRISM dataset. Before training, microbial relative abundance features with a mean abundance less than 0.1% across all samples were removed from the analysis. Models were trained using 10-fold cross-validation. In each partition, 90% of the PRISM dataset was used to train the CGAN model.

CGAN models were trained for 30,000 iterations in which 32 random samples were selected at each iteration as real samples. A synthetic sample was generated for each of the 32 real samples using the sample’s respective disease state as the side information. The 32 real and 32 synthetic samples were then fed to the discriminator for training and the discriminator was updated based on Eq. 1. After updating the discriminator, the discriminator is again used to predict the synthetic samples and the generator is updated based on Eq. 2. Both networks were trained using the ADAM optimizer with a learning rate of 5×10^−5^ (Kingma & Ba, 2014). For the implementation and training of our CGAN models we used the *TensorFlow* package in Python (Abadi et al., 2016).

During training, models were saved every 500 iterations. Additionally, the Principal Coordinate Analysis (PCOA) (Wold, Esbensen, & Geladi, 1987) of the training set, generated set, and the combination of the two sets was visualized and stored. The Bray-Curtis dissimilarity measure was used in calculating the distance matrix for PCOA (Bray & Curtis, 1957). Bray-Curtis dissimilarity quantifies the microbial compositional dissimilarity between two different samples. Given two microbial samples, *x*_*a*_ and *x*_*b*_, the Bray-Curtis dissimilarity between the two samples is calculated as

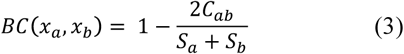

where *C*_*ab*_ is the sum of the lesser values for the abundances of each species found in both *x*_*a*_ and *x*_*b*_, and *S*_*a*_ and *S*_*b*_ are the total number of species counted in *x*_*a*_ and *x*_*b*_ respectively. Visual analysis of the PCOA plots and the overlap of the original and generated data was used to select the best model. An example showing the PCOA of a selected model from the cross-validated training is shown in Fig. 2.

**Figure 2:**
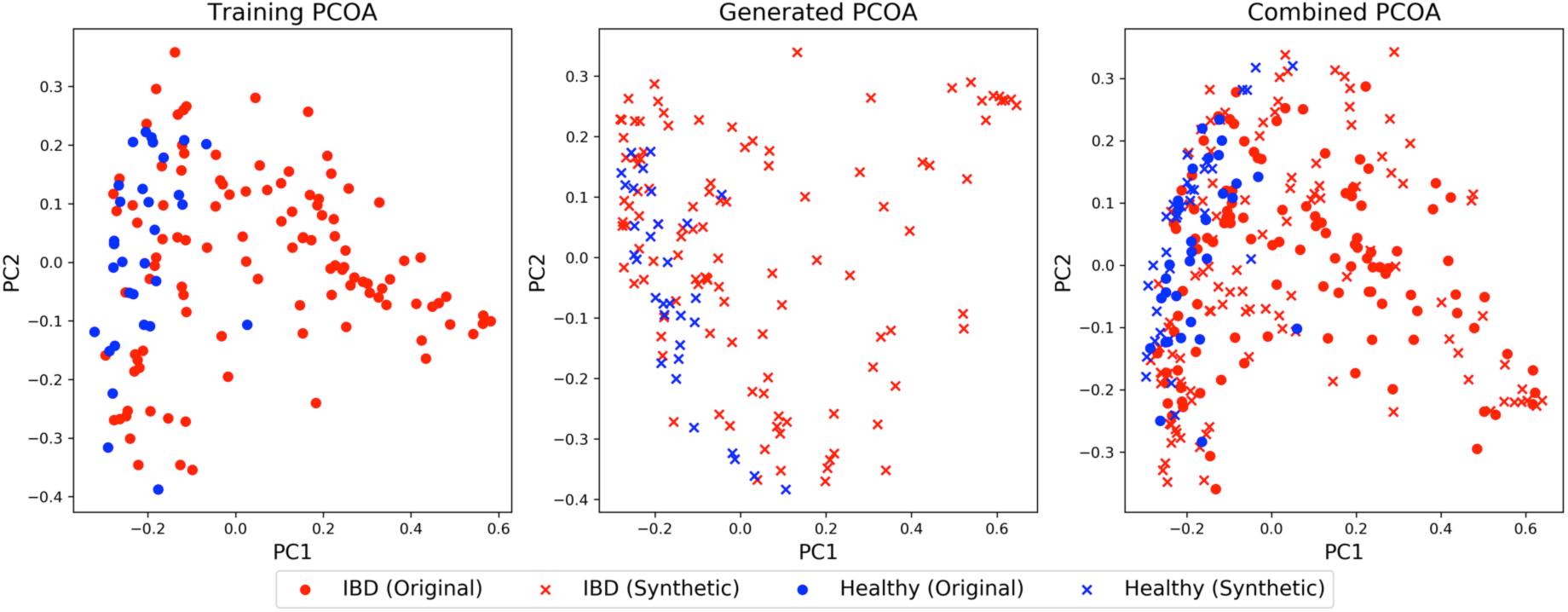
PCOA of the training (left), generated (middles), and combined (right) datasets using Bray-Curtis dissimilarity. Red points represent patients with IBD and blue points represent healthy subjects.

### 3.2 Generated data improves prediction performance

For each of the cross-validation partitions, we simulated 10,000 samples for both IBD and healthy groups using the selected best model. Relative abundance values were then log-transformed and normalized to zero mean and unit variance. Next we trained logistic regression and multilayer perceptron neural network (MLPNN) models on the the fold’s respective training set as well as on the fold’s generated set.

To train the logistic regression model, we performed 5-fold cross-validation grid search over L1, L2, and Elastic Net regularizations considering 10 penalty strengths spaced evenly on a log scale ranging from 1 to 10,000. Logistic regression models were trained using the Python *scikit-learn* package (Pedregosa et al., 2011).

MLPNN models were trained using two fully connected hidden layers with 256 nodes each and dropout with a rate of 0.5 after each layer. Leaky ReLU with an alpha of 0.1 was used as the activation function. The output layer contained two nodes using the softmax activation to predict the disease state. Networks were trained using the ADAM optimizer with a learning rate of 1×10^−4^. We set aside 20% of the training set as a validation set and networks were trained until the loss of the validation set had not decreased for 100 epochs. The implementation and training of the MLPNN models was again done using the *TensorFlow* package in Python (Abadi et al., 2016).

Using the trained logistic regression and MLPNN models generated from a fold’s training set as well as the generated dataset, we calculated the area under the receiver operating characteristic curve (ROC AUC) using the fold’s 10% held out data. We observed that for logistic regression, the models trained using the generated sets had an average ROC AUC of 0.849 while the models trained on the original data had an average ROC AUC of 0.778 across the 10 folds. Similarly, for MLPNN models, the ROC AUC had a value of 0.889 when training on the generated data and 0.847 when training on the original data. Using a Wilcoxon Signed-Rank test, the ROC AUC when using the generated samples was significantly larger than that of when using the original data with a p-value of 0.0249 for logistic regression models and a p-value of 0.0464 for MLPNN models. Boxplots of the ROC AUC values when using original and generated datasets is shown in Fig. 3. These results demonstrated that the CGAN augmented datasets can boost the predictive power of the ML models.

**Figure 3:**
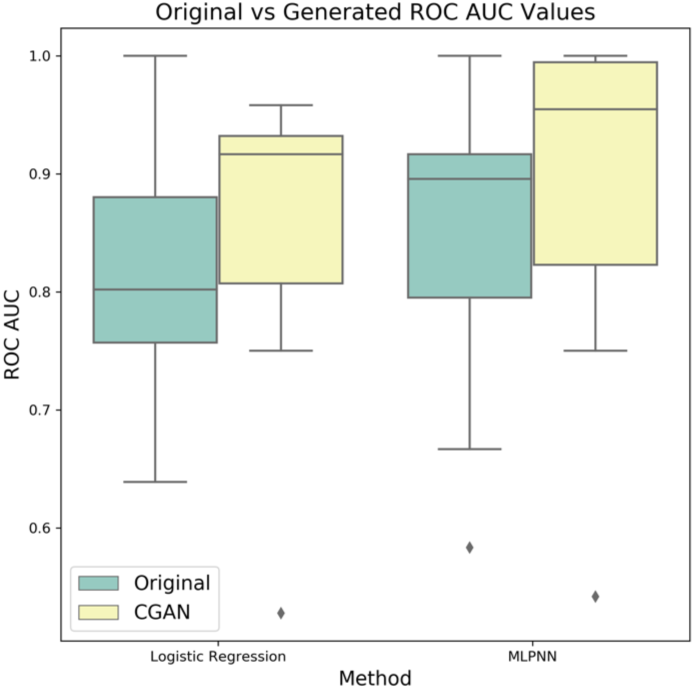
Boxplots for the ROC AUC values across 10-fold cross-validation for logistic regression and MLPNN models hen trained on original and synthetic data.

### 3.3 Diversity of generated data

Diversity metrics are often used to characterize microbiome samples and datasets. In order to check how well the generated distributions of the alpha and beta diversities for IBD and healthy samples. To demonstrate the behaviour of the CGAN model, we visualize the diversity metrics for the training set and for 10,000 generated samples using the selected best model. In addition, we calculate the diversity metrics of a set of 10,000 generated samples using the random initialization of the CGAN before any training to show the initial random distribution.

Alpha diversity is a local measure of species diversity within a sample. It characterizes the microbial richness of a community. For our analysis, we use the Shannon Entropy metric to quantify the alpha diversity of samples. Given a sample *x* with *m* relative abundance values, the Shannon Entropy is calculated as

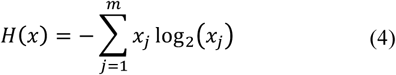

Beta diversity on the other hand considers the ratio between regional and local species diversity and allows us to quantify how similar samples are to each other. In our study, we use the Bray-Curtis dissimilarity as a distance measure of beta diversity, calculated as described in Eq. 3.

Before calculating the diversity metrics, we clipped the generated samples in order to introduce zero values. The softmax function used to generate samples provides a vector entirely of positive values. However, in reality microbiome data is very sparse. Therefore, to induce this sparsity into the generated samples, we calculated the minimum value across all species found in the training set. We used this value as a threshold and set any generated value less than the observed minimum to zero.

After clipping the generated sets, we calculated the diversity metrics. When considering beta diversity, we only considered the Bray-Curtis dissimilarity from the training set to itself, the training set to the best generated samples, and the training set to the randomly generated samples. The distributions of alpha and beta diversity for one of the cross-validated partitions is shown in Fig. 4.

**Figure 4:**
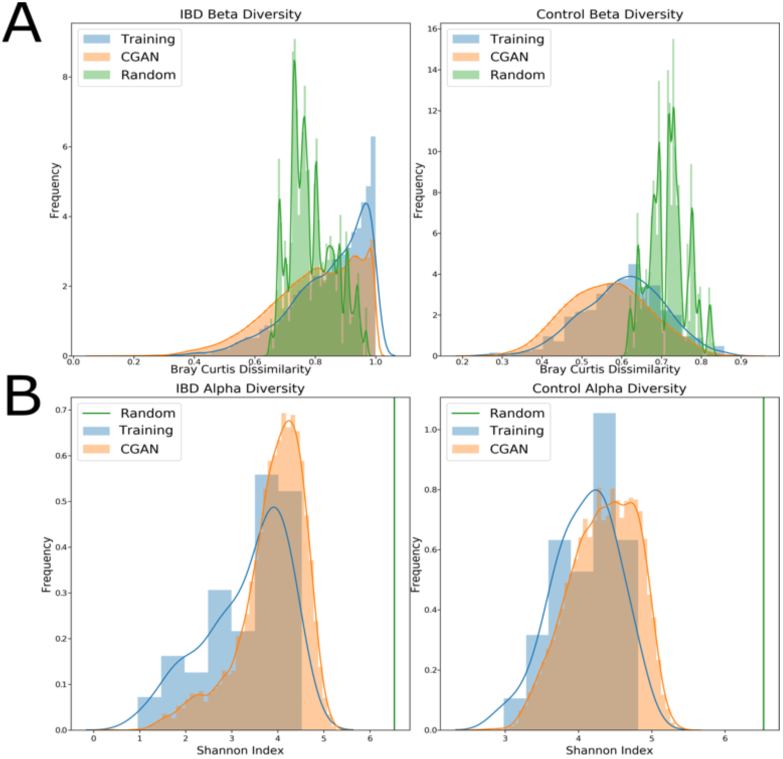
Distributions of the (A) beta diversity based on Bray-Curtis dissimilarity between the training set and itself, the generated (CGAN), and random datasets, and (B) Shannon alpha diversity of training, generated, and random samples for IBD (left) and healthy (right) samples.

We observed that the data generated from the selected best model followed very similar distributions of the alpha and beta diversities of the data used to train the CGAN. We did notice that the beta diversity within the training set had a spike near one, however upon post-analysis we discovered that was caused by samples with only a few number of microbial species present.

### 3.3 Generated data is predictive of external dataset

To evaluate if the synthetic samples generated from the CGAN model were generalizable to a dataset of a similar study, we trained a CGAN model using the entire PRISM dataset in the same manner as described in Section 3.1. The CGAN is trained for 30,000 iterations and models as well as PCOA visualization of the real and synthetic samples are saved every 500 iterations. The best model is selected based on the PCOA comparison between the training and generated sets. A PCOA visualization of the PRISM dataset combined with the synthetic data generated from the best model and the external validation set is shown in Fig. 5

**Figure 5:**
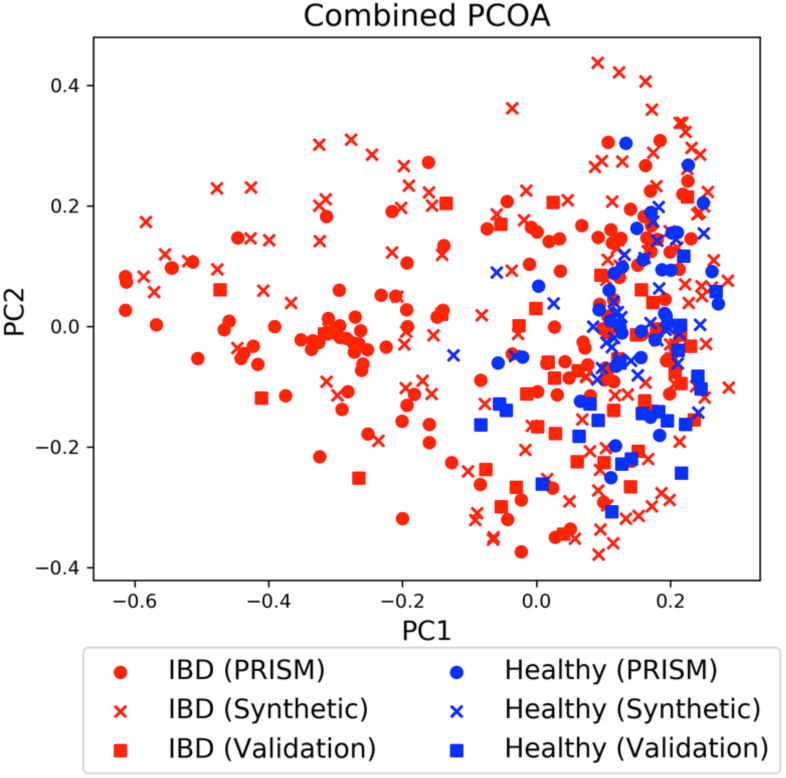
PCOA visualization of the combination of the PRISM dataset, synthetic data generated by the best CGAN model, and the external validation set. Red points represent patients with IBD and blue points represent healthy patients.

Using the best model, we evaluate if generated samples can improve the task of predicting IBD health status. Logistic regression and MLPNN models are trained in the same fashion as outlined in Section 3.2. The model was trained using 10,000 generated samples from a CGAN model trained on the entire PRISM dataset. We then evaluate the model performance on the external validation IBD dataset and observed an improvement in ROC AUC from 0.734 to 0.832 in logistic regression models and fro 0.794 to 0.849 in MLPNN models. This demonstrates that the synthetic samples generated using one cohort are able to augment the analysis of a different cohort.

Lastly, we analyse the distribution of alpha and beta diversities of the original PRISM dataset, the samples generated after training a CGAN on the whole PRISM dataset, and the external validation dataset. Alpha diversity is calculated for each dataset using the Shannon Entropy metric. The beta diversity within the PRISM dataset, from the PRISM dataset to the generated samples, and from the external validation dataset to the generated samples was calculated. In addition, we compared the random diversities from the randomly initialized CGAN before training. Alpha and beta diversities are shown in Fig. 6.

**Figure 6:**
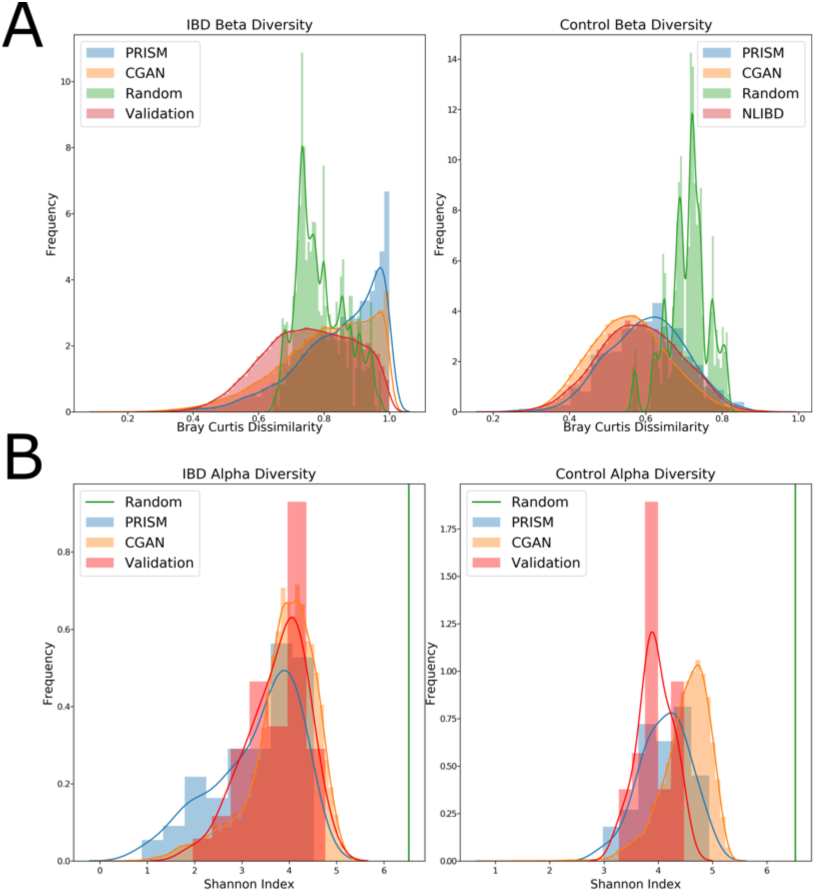
Distributions of the (A) beta diversity based on Bray Curtis dissimilarity between the training set and itself, the validation, the generated (CGAN), and random datasets and (B) Shannon alpha diversity of training, validation, generated, and random samples for IBD (left) and healthy (right) samples.

We observed that the beta-diversity between the PRISM dataset and the synthetic samples generated from it displays similar distributions. Additionally, the distribution of the beta-diversity values between the external validation set and the synthetic samples follow a similar pattern, suggesting that the CGAN model did not overfit the PRISM dataset and is robust in generating synthetic samples. We also observed that the alpha diversities within the PRISM, synthetic, and external validation datasets showed similar distributions. In particular, the alpha diversity within the samples of IBD patients was very similar. The distributions in the healthy samples were slightly different in each of the datasets, however we suspect this may be due to the fact that there were far fewer cases of healthy samples in the original PRISM dataset.

## 4 CONCLUSIONS

In this study, we have developed a novel approach for the generation of synthetic microbiome samples using a CGAN architecture in order to augment ML analyses. Using two different cohorts of subjects with IBD, we have demonstrated that the synthetic samples generated from the CGAN are similar to the original data in both alpha and beta diversity metrics. In addition, we have shown that augmenting the training set by using a large number of synthetic samples can improve the performance of logistic regression and MLPNN in predicting host phenotype.

A current limitation to this approach involves selecting the best CGAN model. Even though visual inspection has been a common approach, it is a subjective and may miss the optimal model. We plan to further this study by investigating stopping criteria using alpha and beta diversity metrics in order to facilitate CGAN model selection.

## Notes

### Competing Interest Statement

The authors have declared no competing interest.

### Summary of Updates

Resubmission included missing equations in the previous version.

